# Prognostic Relevance of CCDC88C (Daple) Transcripts in the Peripheral Blood of Patients with Malignant Melanoma

**DOI:** 10.1101/346551

**Authors:** Ying Dunkel, Anna L. Reid, Jason Ear, Nicolas Aznar, Michael Millward, Elin Gray, Robert Pearce, Melanie Ziman, Pradipta Ghosh

## Abstract

A loss of balance between G-protein activation and deactivation has been implicated in the initiation of melanomas, and non-canonical Wnt signaling via the Wnt5A/Frizzled (FZD) pathway has been shown to be critical for the switch to an invasive phenotype. Daple [CCDC88C gene], a cytosolic guanine nucleotide exchange factor (GEF) which enhances non-canonical Wnt5A/FZD signaling via activation of trimeric G protein, Gai has been shown to serve opposing roles-- as an inducer of EMT and invasiveness and a potent tumor suppressor -- via two isoforms, V1 (full-length) and V2, respectively. Here we report that the relative abundance of these isoforms in the peripheral circulation, presumably largely from circulating tumor cells (CTCs), is a prognostic marker of cutaneous melanomas. Expression of V1 is increased in both the early and late clinical stages (p<0.001, p=0.002, respectively); V2 is decreased exclusively in the late clinical stage (p=0.011). The two isoforms have opposing prognostic effects: high expression of V2 increases progression-free survival (PFS; *p* = 0.02), whereas high expression of V1 decreases PFS (p=0.013). Furthermore, these effects are additive, in that melanoma patients with a low V2-high V1 signature carry the highest risk of metastatic disease. We conclude that detection of Daple transcripts in the peripheral blood (i.e., liquid biopsies) of patients with melanoma may serve as a prognostic marker and an effective strategy for non-invasive long-term follow-up of patients with melanoma.

## Introduction

Melanoma is a highly malignant cancer and causes more deaths than other forms of skin cancer; as in most cancers, early detection and surgical intervention are imperative for a curative outcome. However, in the case of melanoma, even with ‘curative’ surgical resection, the risk of recurrence is never completely absent and late recurrences, even beyond 8 y, can occur at a frequency ranging from 0.41% to 25% across different studies, with 6.8% at 15 y and 11.3% at 25 y ^1,2^. The late recurrence population is particularly interesting to study as it represents a group of patients exhibiting tumor dormancy, defined as a stage in cancer progression in which residual disease is present but remains asymptomatic ^2^. Hence, early detection of patients in this clinical stage of melanoma is of paramount importance so that they can be offered adjuvant therapy.

Tumor dormancy is best studied by examining patients who demonstrate persistent circulating or disseminated tumor cells (CTC or DTC) after removal of all clinically evident disease ^3^. In fact, metastatic melanoma was the first malignancy in which CTCs were detected^4^. More recently, the presence of circulating melanoma cells (CMCs) in the peripheral blood of patients with metastatic cutaneous melanoma has been identified by detecting mRNA transcripts of specific melanoma markers. Transcripts from one or more circulating melanoma cells (CMCs) in 10 ml of peripheral blood can be identified in 45% of all melanoma patients at varying disease stages (AJCC Stages I /II, III, and IV: 35%, 44%, and 86%, respectively) ^5–9^. While the prognostic impact of the number of CMCs remains to be determined, the prognostic impact of detecting molecular markers from these CMCs in the peripheral blood of melanoma patients has been evaluated by multiple groups within the last decade. These studies are in general agreement that the detection of unique multi-panel biomarkers such as S100B, MAGE-3, p97, MUC-18, S100A4, MART1, HMB45, and tyrosinase, by real-time quantitative PCR (RT-PCR) in CMCs of melanoma patients can have a valuable prognostic utility, especially in patients with Stage II and III disease ^10–14^. These studies demonstrate that molecular detection of CMCs may be valuable to detect early disease recurrence and to stratify patients for adjuvant therapy. However, given the heterogeneous nature of melanoma cells, no single marker or panel of markers appears to be sufficiently robust in predicting outcome.

Despite these insights, the identification of CMC-biomarkers with strong rationale and mechanistic insights and specifically within signaling pathways known to fuel melanoma metastasis, remains unexplored and unrealized. In this regard, several studies agree that activation of non-canonical Wnt signaling and inactivation of the canonical Wnt pathway is associated with metastatic features of melanoma and thus poor survival ^15–17^. Here we report the identification of Daple (CCDC88C), a cytoplasmic transducer of Wnt signals ^18–20^, as a candidate marker for monitoring deregulated Wnt signaling during melanoma progression. Daple links ligand-activated Frizzled receptors to Gi protein activation^18^. Such activation, on one hand antagonizes the canonical Wnt pathway and suppresses a neoplastic transformation and tumor growth. On the other hand, it enhances non-canonical Wnt pathway and Akt signals, triggering epithelial-mesenchymal transition (EMT) and tumor invasion ^18^. Subsequent work unraveled that Daple-dependent modulation of Wnt signaling is triggered not just by Wnt ligands ^18^, but also by growth factors (EGF) ^20^ and junctions-associated Cadherin-catenin complexes that trigger the PI3K-Akt pathway ^19^. Perhaps as a consequence of such convergent signaling, ourselves and others have demonstrated Daple’s role in fueling the metastatic progression of gastric ^21^ and colorectal cancers (CRCs) ^18,22^ and its ability to serve as a prognostic marker in the CTCs of patients with metastatic CRCs ^22,23^. We hypothesized that detection of Daple transcripts in the peripheral blood of patients with melanoma may serve as a surrogate marker for deregulated Wnt signaling during melanoma progression. Our findings present evidence that tracking levels of its transcripts could serve as an effective strategy for non-invasive long-term follow-up of patients with melanoma.

## Results and Discussion

### Rationale for choosing Daple as a biomarker for metastatic progression of melanoma

We previously reported that the Disheveled-binding protein, Daple, is a guanine nucleotide <u>e</u>xchange <u>m</u>odulator (GEM) for heterotrimeric G proteins within the Wnt signaling cascade ^18^. Subsequent work revealed that the human Daple/CCDC88C gene is expressed as two functional isoforms: Daple-full length (V1) (2028aa) and Daple-V2 (552aa) ^22^. These isoforms serve contrasting roles during cancer progression-- while both isoforms cooperatively antagonize Wnt signaling and suppress tumor cell proliferation and growth via their GBA motif, only the full-length V1 isoform triggers EMT and invasion. In the colon, both isoforms collaboratively suppress tumor cell growth, and have an additive prognostic impact (worse outcome when both are suppressed). However, only Daple-V1 is increased in invasive tumor margins and in CTCs disseminated during CRCs progression. We next chose to study the role of Daple in melanoma, which, like colorectal cancers, is another cancer that is characterized by dysregulation of heterotrimeric G-protein as well as Wnt signaling; prior studies have implicated both pathways in the metastatic progression of melanomas ^24,25^. When we analyzed the RNA sequencing dataset from The Cancer Genome Atlas (TCGA) for dysregulation of Daple/CCDC88C expression (CNVs, deletions or mutations) and the frequently mutated heterotrimeric G-protein GNAQ^26^, we found that mutations have been identified in patients with cutaneous, but not uveal melanomas for Daple/CCDC88C. For GNAQ, on the other hand, although mutations have been identified in both cutaneous and uveal melanomas, these are predominantly found in uveal melanomas (**Fig 1A-B**). Mutations found in Daple/CCDC88C included both missense and truncating mutations (**Supplemental Fig S1-S2**). Notably, mutations in other heterotrimeric G-protein α-subunits (GNAI1, GNAI2, GNAI3, and GNAS) that are modulated by this family of GEMs (of which Daple is a member and CCDC88A/Girdin and NUCB1/2 are others) were only found in cutaneous melanomas (**Fig 1B**). These findings, along with the observation that Daple/CCDC88C was indeed abundantly expressed in distant metastases to diverse organs or sites (**Supplemental Fig S3**), led us to hypothesize that dysregulated signaling via Daple/CCDC88C in cutaneous melanomas may have a role in the progression to metastatic disease.

**Figure 1.**
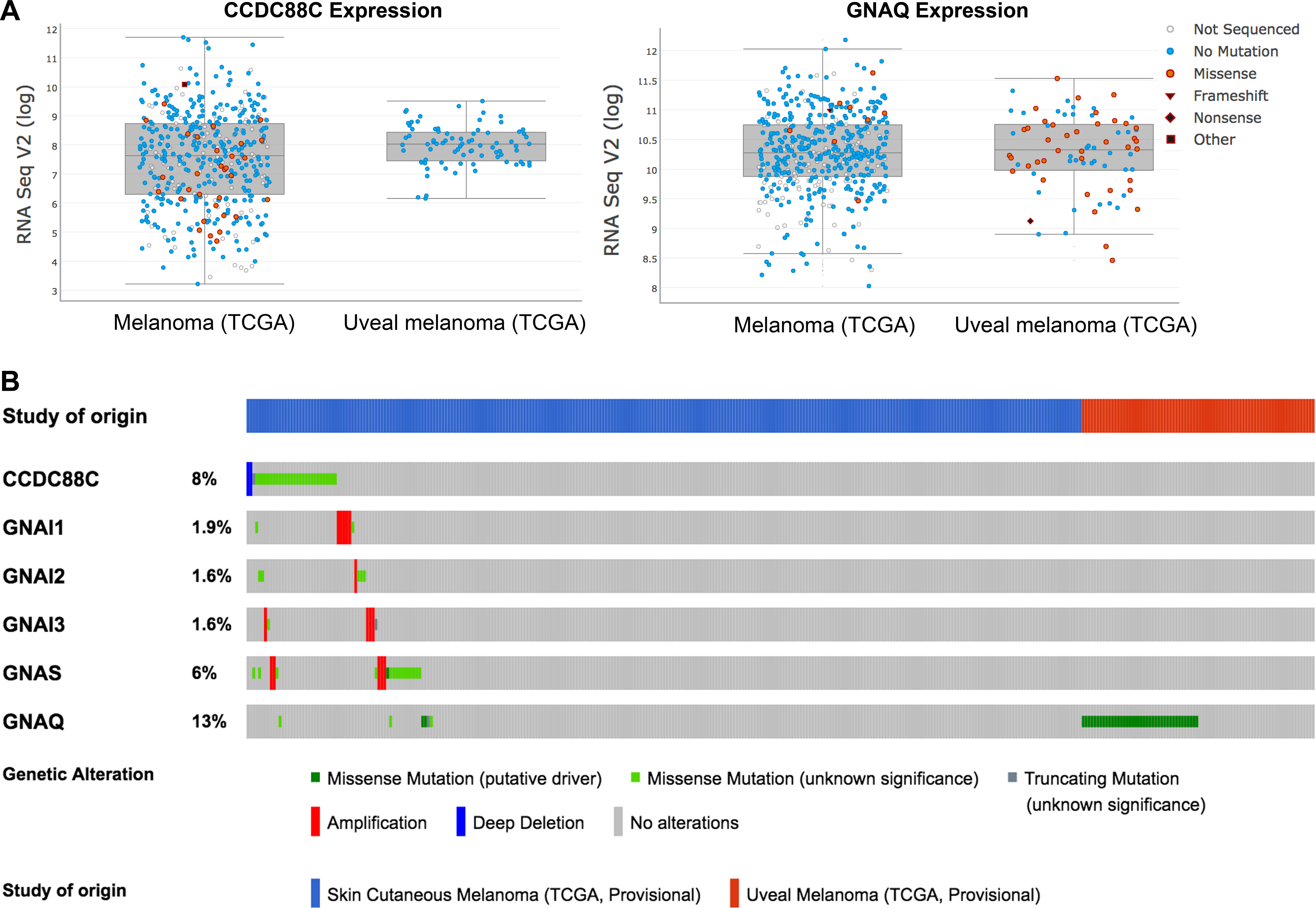
Daple expression is altered (deleted or mutated) in cutaneous, but not in uveal melanomas. (**A**) Graphs display the levels of expression of CCDC88C/Daple (*left*) and GNAQ (*right*) in cutaneous melanoma vs. uveal melanoma patients in The Cancer Genome Atlas (TCGA) dataset. Individual point represents mutational status of each patient. While CCDC88C mutations are restricted to cutaneous melanomas, GNAQ mutations are found predominantly in uveal melanomas. (**B**) Frequency of gene alterations in cutaneous melanoma vs. uveal melanoma patients in The Cancer Genome Atlas (TCGA) dataset is shown for CCDC88C (Daple) and heterotrimeric G-proteins (GNAI1, GNAI2, GNAI3, GNAS, and GNAQ). Mutations in CCDC88C and GNAI and GNAS proteins were observed exclusively in cutaneous melanomas. Mutations in GNAQ was predominantly observed in uveal melanomas.

### Transcripts of Daple-V1 (V1), but not Daple-V2, is increased in the peripheral circulation of patients with melanoma

We recently showed that the expression of Daple full-length (henceforth, Daple-V1) was elevated in EpCAM-immuno-isolated CTCs of colorectal cancer patients, and that Daple promotes epithelial-to-mesenchymal transition (EMT) during progression ^18,22^. Expression of Daple-V2, on the other hand, was found to be decreased in primary CRC tumors^22^. Because Daple-V1 and-V2 are differentially expressed in CTCs of CRCs patients we first asked if this is also the case in patients with melanoma (see Table 1 for patient cohort characteristics). Because detection of melanoma transcripts in the peripheral blood is a sensitive surrogate marker of melanoma CTCs^8,27^, we analyzed the abundance of Daple transcripts in the peripheral blood of patients with melanoma (n = 205) and of healthy controls (n = 142) by qRT-PCR. We found that patients with melanoma have significantly elevated levels of Daple-V1 expression compared to normal healthy controls (*p* < 0.0001; **Fig. 2A, B**). The same pattern of expression was observed as for the melanocyte marker and the metastasis-associated protein, S100A4 ^28^, which is a transcriptional target of the Wnt-β-Catenin pathway^29^ and currently used as a marker to diagnose melanoma^30^. By contrast, no difference in Daple-V2 expression was observed between transcripts in peripheral blood of melanoma patients and the healthy controls (**Fig 2A-B**). These findings indicate that transcripts of full length Daple-V1, but not V2 are selectively upregulated in patients with melanoma and suggest that high levels of Daple-V1 may contribute to melanoma progression.

**Figure 2.**
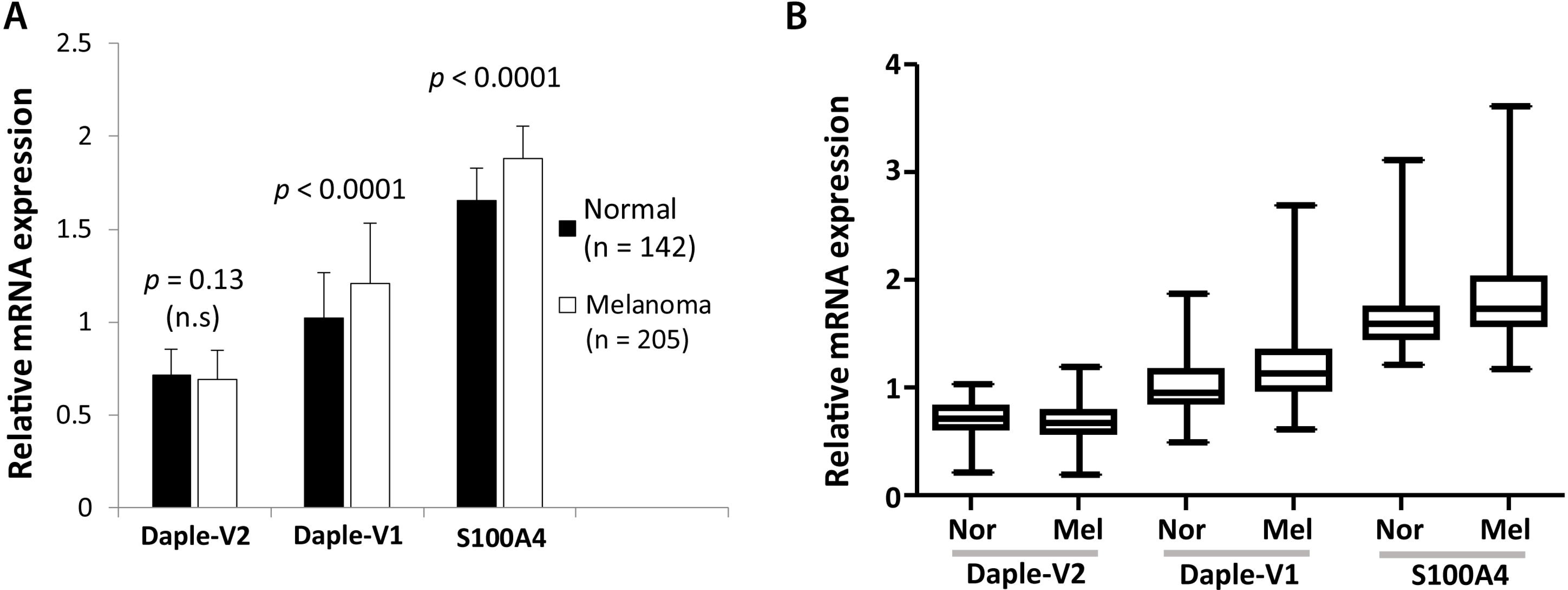
Comparison of the expression of Daple (isoforms V1 and V2) and S100A4 in the peripheral blood of patients with cutaneous melanoma *vs.* normal healthy subjects. Expression of Daple V1, V2 and S100A4 (positive control) mRNA were analyzed in patients with melanoma (n = 205) and in normal healthy controls (n = 142) by qPCR. Findings are displayed as bar graphs (**A**) and box plots (**B**) for the relative mRNA expression of each gene normalized to GAPDH using standard curve method. The statistical significance of the difference for each gene between the melanoma group and the normal control group was calculated by t-test. Levels of expression of Daple-V1 and S100A4 (positive control), but not Daple-V2 were significantly elevated in patients with melanoma compared to healthy normal subjects.

### Transcripts of Daple-V1 increase, but Daple-V2 decrease in the peripheral blood of patients during melanoma progression

To further dissect the patterns of expression of Daple-V1 and Daple-V2 relative to disease stage, we analyzed both isoforms in the peripheral circulation of patients with early stage (AJCC stages 0, I, II) vs. late stage (AJCC stages III, IV) melanomas. We found that Daple-V1 expression is significantly increased in both early (*p* < 0.001; **Fig. 3A**) and late (*p* = 0.002; **Fig. 3A**) stages compared to healthy controls. Expression of Daple-V2 is significantly lower in late stage melanoma compared to early stage disease and healthy controls (*p* = 0.008 and 0.003, respectively; **Fig. 3B**); no significant differences were noted between healthy and early stage melanoma patients. Levels of expression of S100A4, the positive control we used in this study, was increased in both early (0, I, II) and late (III, IV) stages (**Fig. 3C**) compared to normal healthy controls (*p* < 0.001 and 0.01, respectively). The changes in Daple expression we observe relative to disease stage are consistent with the patterns of Daple-V1 vs. V2 expression in CRC stages; in both scenarios Daple-V1 increases but Daple-V2 decreases as the disease progresses from the early state to the late invasive stage and undergoes hematogenous dissemination. The change in these transcripts in peripheral blood may reflect the entrance of CTCs into the peripheral circulation ^22^.

**Figure 3.**
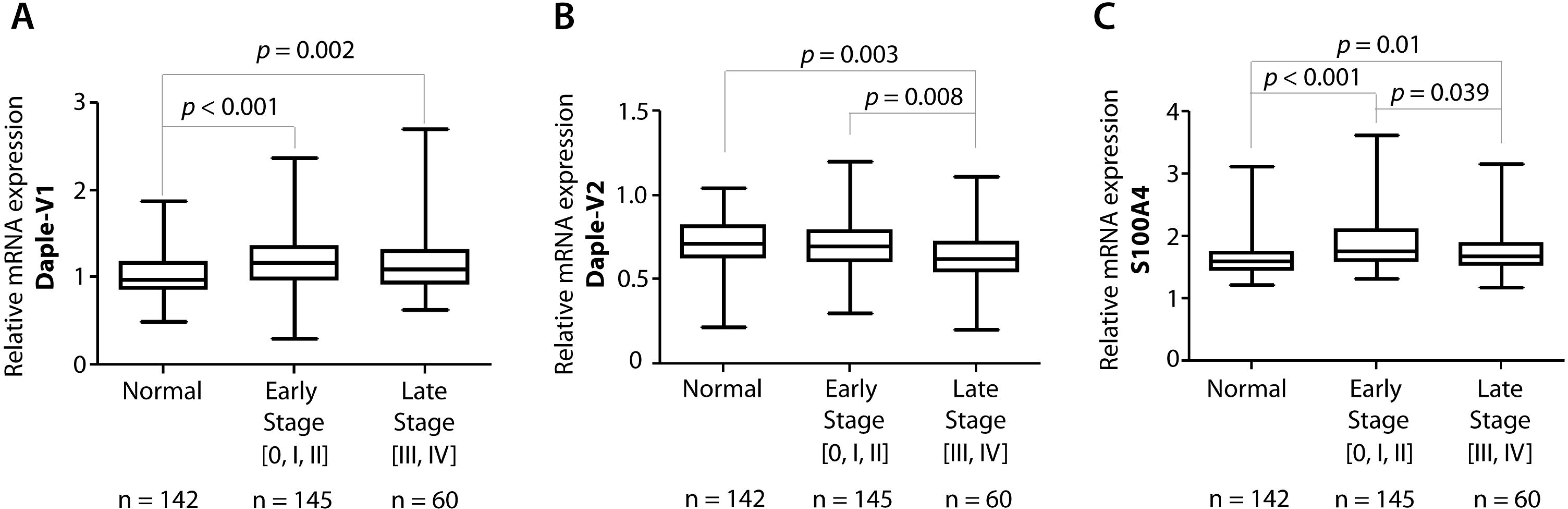
The expression of Daple isoforms V1 (A) and V2 (B) and S100A4 (C) in the peripheral blood of normal subjects and of patients with early or late-stage melanomas. Expression of Daple V1, V2 and S100A4 (positive control) mRNA were analyzed in patients with early (clinical stage 0, I, and II; n = 145) or late (clinical stage III and IV; n = 60) stage melanoma and in normal healthy controls (n = 142) by qPCR. Findings are displayed as box plots for the relative mRNA expression of each gene normalized to GAPDH as in **Figure 2**. The statistical significance of the difference for each gene between groups with either early or late -stage melanoma and normal healthy controls was calculated by t-test. While transcripts of Daple-V1 (**A**) and S100A4 (**C**) are elevated in both early and late stage melanomas, those of Daple-V2 is reduced exclusively in late stages of melanoma.

### Elevated Daple-V1 and suppressed Daple-V2 transcripts in the peripheral circulation of patients with early stage melanoma carries an increased risk for distant metastasis

Next, we asked if the levels of expression of Daple-V1 and Daple-V2 in the peripheral blood of early stage melanoma patients can correlate with the frequency of progression to metastatic disease. In 145 patients with early stage melanoma, who progressed to metastatic disease, we found that low expression of Daple-V2 was associated with a higher metastatic rate (14.44%) compared to high expression of Daple-V2 (1.82%; *p* = 0.0174; **Fig. 4A**). This pattern was not seen in the case of Daple-V1; in fact, high expression of Daple-V1 tracked with higher metastatic rate (16.33%) compared to low expression of Daple-V1, although that trend did not reach significance (6.25%; *p* = 0.073; **Fig. 4B**). When we compared the rates of metastatic progression between a group of patients with a high Daple-V1-low Daple-V2 signature and another group with low Daple-V1-high Daple-V2 signature, we found that the former has a much higher metastatic rate (36.36%) compared to the latter (3.57%; *p* = 0.006; **Fig. 4C**). These findings suggest that an analysis that accounts for the levels of expression of both Daple isoforms could perform better in prognosticating outcome than each one alone, i.e., Daple-V1 and Daple-V2 may have an additive prognostic impact. The high expression of the metastasis-associated protein, S100A4^28^ was also somewhat indicative of a higher metastatic rate among the high expressing group (13.41%) compared to the low expressing group (4.76%; *p* = 0.095; **Fig. 4D**) but did not reach significance. We speculate that the prometastatic trends we observed in the case of both Daple-V1 and S100A4 did not reach significance because of insufficient patients in the cohort. Regardless, these findings are consistent with our recent findings that Daple-V2 is a potent tumor suppressor, whereas Daple-V1 triggers EMT and invasion ^18,22^.

**Figure 4.**
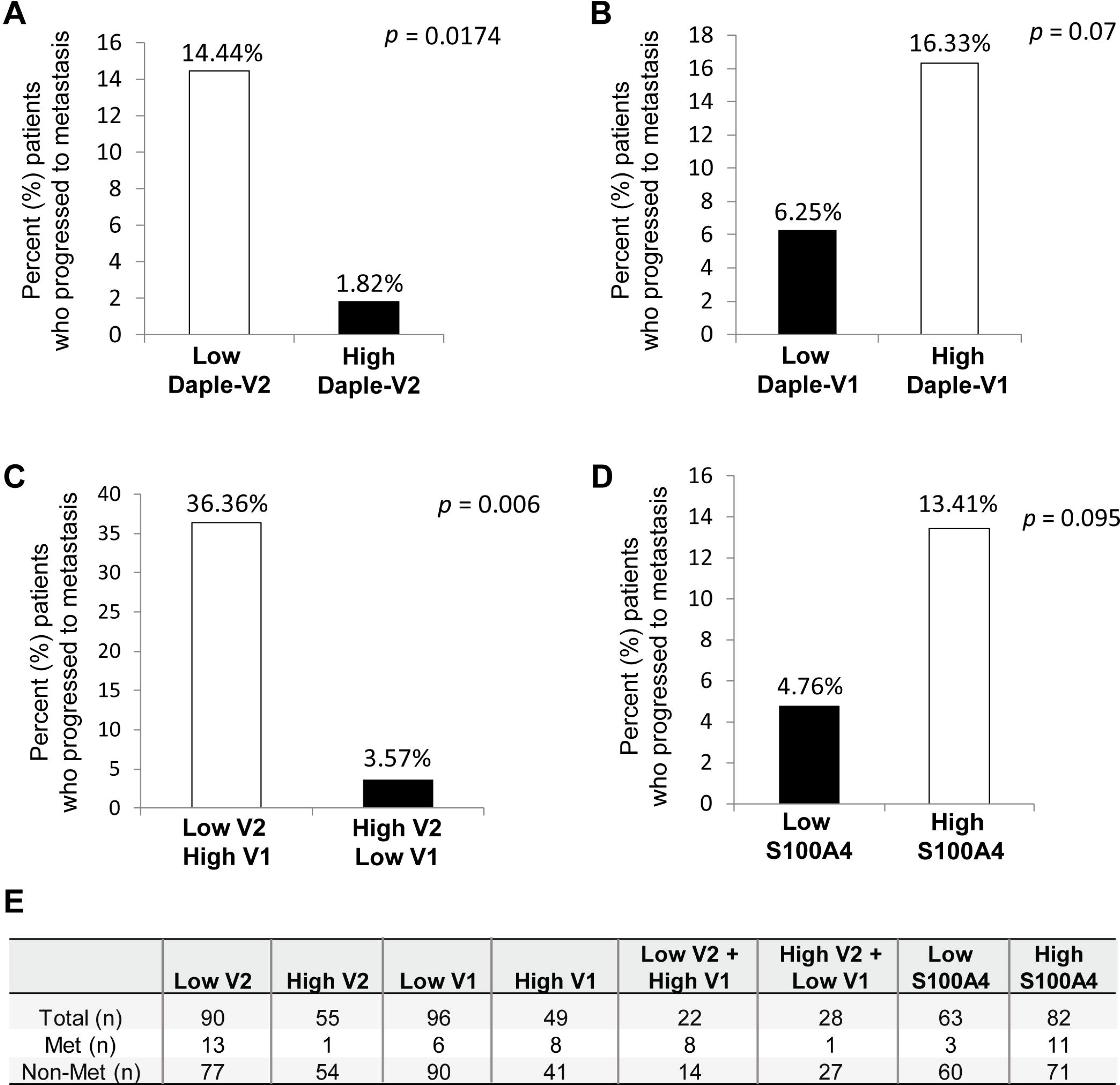
Frequency of progression to metastasis among patients stratified based on high vs. low levels of circulating transcripts of Daple-V2 alone (A), Daple-V1 alone (B), Daple V1 and V2 combined (C), and S100A4 (D) in the peripheral blood. Bar graphs display the proportion of patients who progressed to metastasis (Y axis) in patients with low vs. high expression of indicated genes. (**A**) Compared to those with high Daple-V2 expression (n=55), those with low Daple-V2 (n = 90) expression displayed higher frequency of metastatic progression. (**B**) Compared to those with low Daple-V1 expression (n = 96), those with high Daple-V1 (n = 49) expression displayed higher frequency of metastatic progression. (**C**) Compared to those with low-V1-high-V2 signature (n = 28), those with high-V1-low-V2 (n = 22) expression displayed higher frequency of metastatic progression. (**D**) Compared to those with low S100A4 expression (n = 63), those with high S100A4 (n = 82) expression displayed higher frequency of metastatic progression. Optimal cut-off values for each gene expression were statistically derived (see detailed *Methods*) to divide patients into subgroups with low *vs*. high expression levels. Fisher exact test was used to compare the low vs. high subgroups.

Finally, we also found that patients with low Daple-V2 transcripts in their peripheral circulation had higher levels of other melanoma cell markers (**Fig 5A-B**; **Supplementary Table S4**; data from^7^), indicating that these patients may have higher numbers of circulating melanoma cells. Taken together, these findings suggest that Daple-V2 alone may serve as an effective prognostic marker in early stage melanoma.

**Figure 5.**
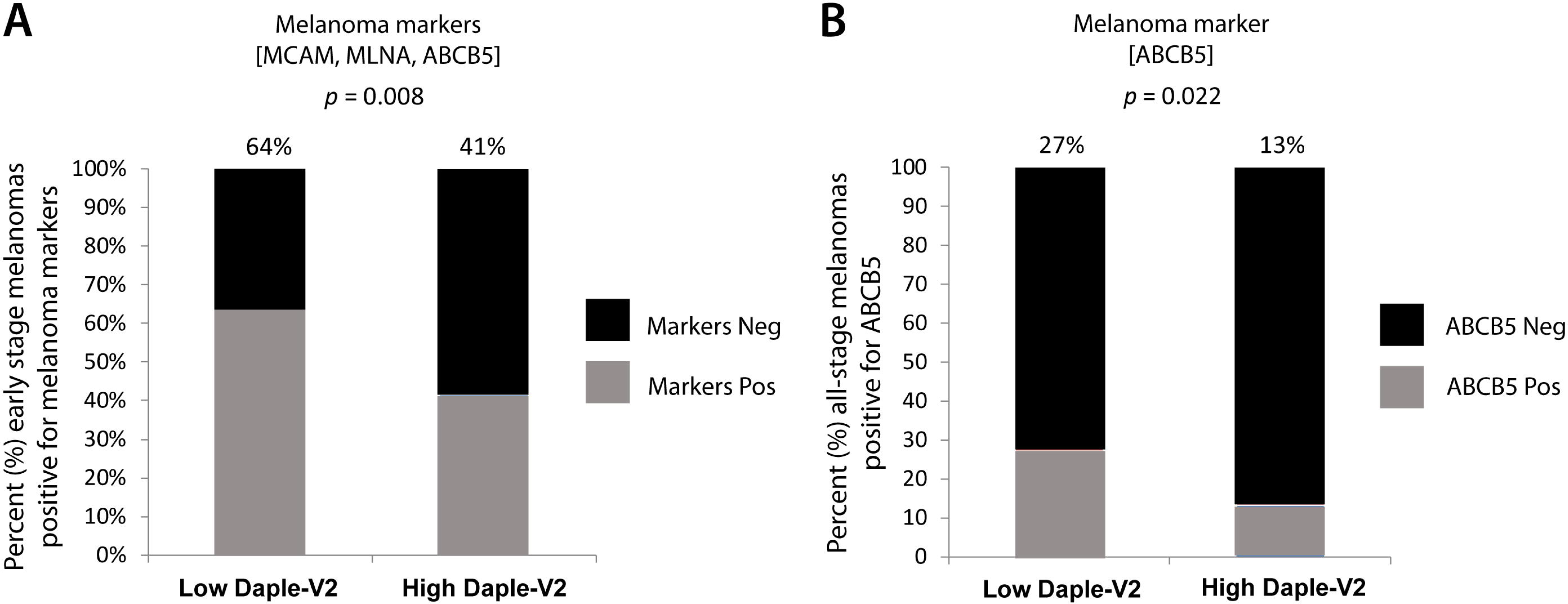
Correlation between Daple-V2 expression and melanoma-associated markers. (A) Patients with early stage melanomas (n = 145) with known expression status (positive or negative) of melanoma associate markers (MCAM, MLNA and ABCB5) were analyzed in subgroups of patients with high (n = 55) or low (n = 90) expression levels for Daple-V2. Tumors were classified as “Markers Pos” if any one of the three melanoma-associated markers were detectable, whereas tumors were classified as “Markers Neg” if none of the three markers were detectable [see **Table S4-source data**]. Bar graphs display the incidence of marker positivity as percent (%; Y axis) of patients with low vs. high Daple-V2. (**B**) Patients with all stages of melanoma (n = 205) with known expression status (positive or negative) of the melanoma stem cell marker, ABCB5 alone was analyzed in subgroup of patients with low vs. high Daple-V2 as in **A**.

### Prognostic impact of Daple-V1 and Daple-V2 in patients with melanoma

Because the frequency of metastatic progression was different for patients with high *vs*. low expression of transcripts for Daple-V1 or-V2 in the peripheral circulation at an early stage of disease (stage 0, I, II), we asked if levels of these transcripts can prognosticate progression free survival (PFS). Kaplan-Meier curves for high vs. low levels of expression of Daple-V1 and -V2 were plotted and compared between the 2 groups using the log rank test. In the case of Daple-V2, we found that PFS for patients with low expression was significantly reduced compared to patients with high expression (p=0.0139; **Fig. 6A**). While in the case of Daple-V1, we found that patients with high expression tend to have shorter PFS compared to patients with low expression (*p* = 0.0833; **Fig. 6B**). A similar pattern was found in PFS analysis of S100A4; patients with high expression tend to have shorter PFS compared to patients with low expression but this did not reach significance (*p* = 0.051; **Fig. 6C**). These results indicate that Daple-V2, but not V1 is a favorable prognostic indicator of PFS. Next we asked if the prognostic impact of Daple-V1 and Daple-V2 are additive. When we compared PFS between the group with low Daple-V1-high Daple-V2 and the group with high Daple-V1-low Daple-V2, we found that patients with low Daple-V1-high Daple-V2 have a much higher PFS than the other group (*p* = 0.013; **Fig. 6D-E**), indicating that Daple-V1 improves the prognostic value of Daple-V2.

**Figure 6.**
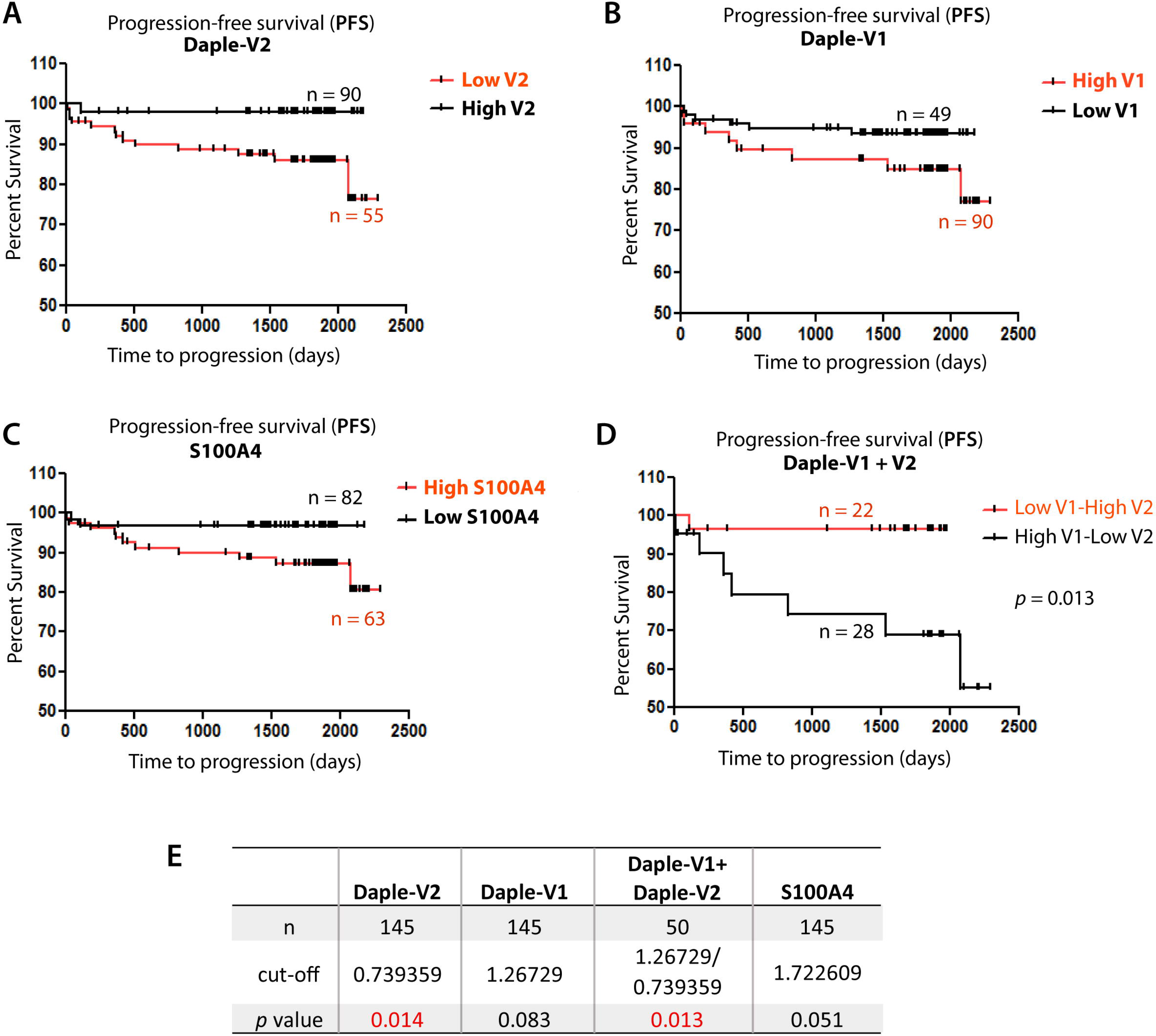
Association of progression-free survival (PFS) with expression level of Daple isoforms V2 and V1 (A, B, D), and S100A4 (C) Kaplan-Meier plots for PFS (% survival; y-axis) of 145 patients with early stage melanoma was stratified based on levels of Daple-V2 alone (**A**), Daple-V1 alone (**B**), S100A4 alone (**C**) and Daple-V1+V2 combined (**D**) and plotted against duration of follow-up (days to progression; x-axis). Although high Daple-V2, low Daple-V1 and low S100A4 carried better prognosis, table (in **E**) shows that statistical significance was reached only for Daple-V2. The positive prognostic impact of Daple-V2 and the negative prognostic impact of Daple-V1 were additive (**D-E**).

### Stratification of patients by accounting for the expression levels of both Daple-V1 and Daple-V2 offers novel prognostic signatures

To identify the most effective diagnostic algorithm that combines the prognostic impacts of both Daple isoforms, we explored if assessment of Daple-V1 among the patients with low Daple-V2 can improve the prognostic impact of Daple-V2. When we stratified patients with low Daple-V2 into two groups with high and low Daple-V1, we found that patients with high expression of Daple-V1 have shorter PFS than those with low expression of Daple-V1 (*p* = 0.0011; **Fig. 7A**). PFS among patients with low Daple-V2-high Daple-V1 expression was significantly lower than those with high Daple-V2 expression (*p* < 0.0001; **Fig. 7B**).

**Figure 7.**
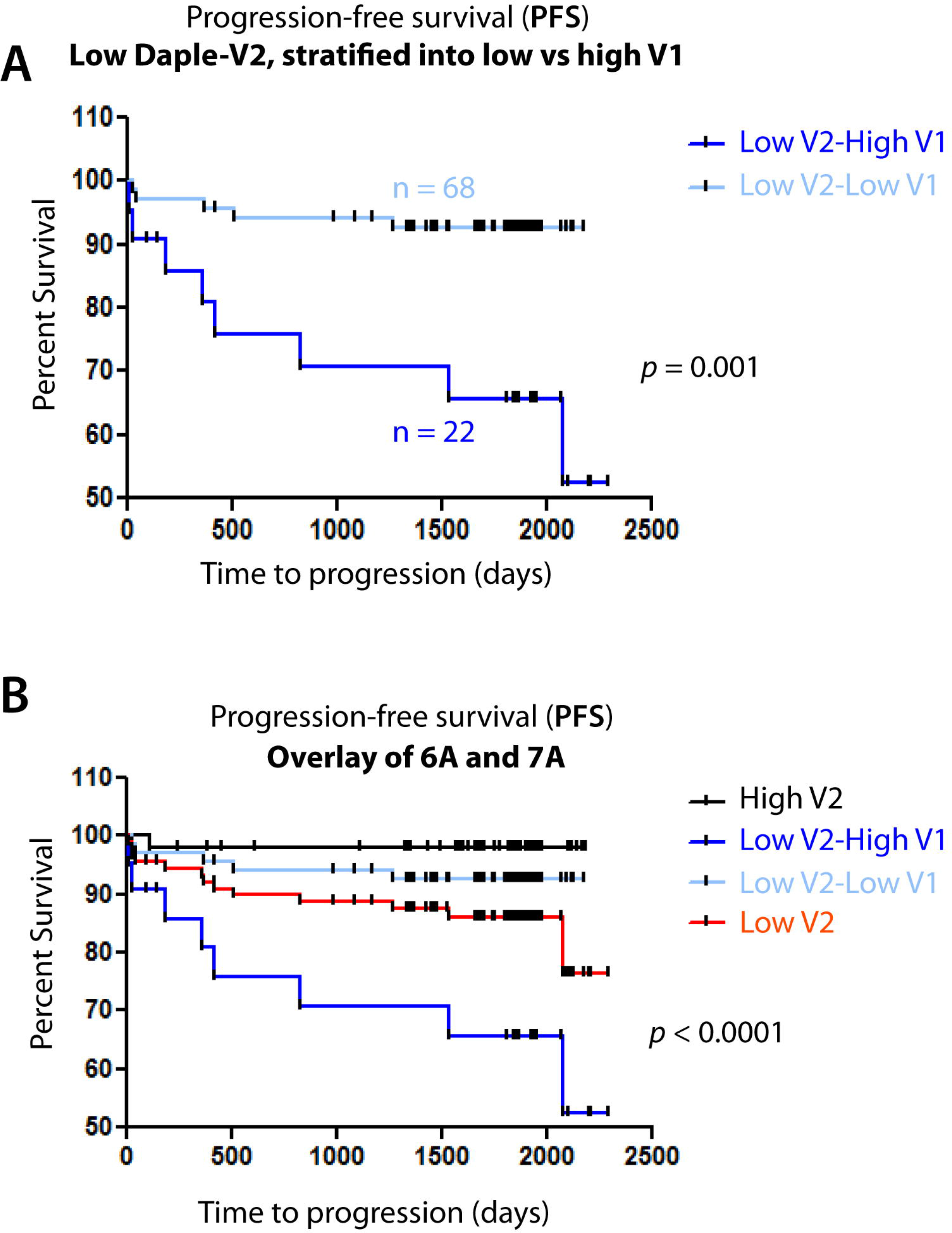
Stratification of patients with low Daple-V2 based on their levels of expression of Daple V1 further improves the prognostic power of Daple-V2. (**A**) Kaplan-Meier plots for PFS (% survival; y-axis) of 90 patients with early stage melanoma with low levels of Daple-V2 expression was stratified based on levels of expression of Daple-V1. (**B**) An overlay of Kaplan-Meier plots displayed in **Fig. 6A** [i.e., PFS among patients stratified based on Daple-V2 alone] and **Fig. 7A**.

We further evaluated the effect of Daple-V1 and -V2 on survival rates after adjustment for clinical covariates including age and tumor stage using Cox proportional hazard regression analysis (Cox analyses). We found that Daple-V2 can be an independent prognostic factor; PFS of patients with low Daple-V2 expression was significantly lower than those with high Daple-V2 (hazard ratio [HR] 9.1059; *p* = 0.00362; 95% confidence interval [CI] 1.1522 to 71.9672). Cox analyses also confirmed that Daple-V1 and Daple-V2 have additive prognostic value; PFS was significantly reduced when patients have high Daple-V1 and low Daple-V2 (HR 13.1368; *p* = 0.0172; CI 1.5792 to 109.2828). Although statistical significance was reached in each case, it is noteworthy that the 95% CIs are wide and further studies with larger cohort sizes are warranted. Regardless, both log rank and Cox analyses are in agreement that Daple-V2 alone is a sufficient prognostic factor, and that risk stratification for tumor progression improves by the additive effect of Daple-V1 and Daple-V2. Risk was lowest when patients have low Daple-V1-high Daple-V2 signature, and higher when patients have a high Daple-V1-low Daple-V2 signature.

## Conclusions

The most important discovery we report here is a unique Daple/CCDC88C transcript signature, ‘high Daple-V1 and low Daple-V2’ in the peripheral blood of patients with early stage cutaneous melanoma that effectively stratifies patients at the highest risk for progression to metastatic disease. Daple-V1 triggers EMT via enhancement of non-canonical (β-catenin-independent) Wnt signals and suppresses canonical (β-catenin-dependent) Wnt signals ^18^, whereas Daple-V2 serves primarily as a potent tumor suppressor ^22^. Therefore, an increase in Daple-V1 at early stages in the peripheral blood (most likely from CMCs), suggests that non-canonical Wnt signals are likely enhanced and canonical Wnt signals are suppressed early during melanoma progression, a signature that has been previously described as a contributor to metastatic features of melanoma and poor survival ^15–17,31^. A decrease in the tumor suppressive isoform, Daple-V2 at late stages suggests that canonical Wnt signals are upregulated and may be necessary for the growth of metastatic lesions at distant sites. Overall, an increase in Daple-V1 and a decrease in Daple-V2 expression during disease progression is indicative of an overall permissive signaling program for melanoma progression, and this notion is consistent with the prognostic signature that emerged from this work as the most effective in predicting the risk for metastatic progression.

It is noteworthy that although lower Daple-V2 transcripts in the peripheral circulation were seen predominantly in patients with late stage disease, lower Daple-V2 transcripts in the early stage of the disease emerged as the strongest predictive profile in our study. This came as an unusual surprise to us because typically biomarkers representative of CTCs are elevated in circulation over background, not decreased. These findings raise the possibility that circulating melanocytes may be present in the peripheral circulation of healthy controls (hence, Daple-V2 transcript is detected in the circulation of normal healthy individuals), and that metastatic progression of melanoma may be associated with dissemination of different populations of CMCs that have low Daple-V2. In fact, this possibility is supported by prior studies that reported that melanocyte precursor cells – cells which express TYR and MITF transcripts, but not MLANA (or MIF) - may indeed circulate in healthy adults with benign nevus, are detected in human cord blood,^32–34^ and during melanocytic differentiation into vein-like structures from pluripotent stem cells ^35^. In patients with melanoma, a shift is noted in the landscape of the melanoma transcriptome, i.e., MLANA and MIF transcripts are elevated (but not TYR or MITF) ^33^, leading some to speculate that the circulation of normal melanocytic precursors is inhibited in patients with melanomas, perhaps due to factors, e.g., cytokines released into the systemic circulation ^33^. Based on these studies, and our own findings, we conclude that the entry of normal melanocytes and/or its precursors that have high Daple-V2 is inhibited in patients with melanoma, and instead, melanoma cells with lower Daple-V2 begin to enter the circulation. Because the burden of metastatic CMCs in the circulation increases with increasing clinical stage of disease ^5^, the prognostic value of decreased Daple-V2 transcript that we observe in the early stage suggests that it may be a sensitive method to detect the changing landscape of melanocyte transcripts and serve as a surrogate marker for the overall load of metastatic CMCs.

Overall, we conclude that abundance of Daple transcripts (V1 and V2) in the peripheral blood of patients with melanoma may serve as a clinically useful early prognostic test for detecting sinister CMCs in the circulation. Because malignant melanoma is a disease that can be characterized by prolonged periods of tumor dormancy, which necessitates prolonged periods of follow-up, monitoring Daple transcripts in the peripheral circulation may serve as a relatively simple minimally-invasive test, either administered stand-alone, or in combination with other panels of melanoma markers. Further studies in larger cohorts are necessary before one can rigorously assess and realize the clinical usefulness of Daple in melanoma surveillance during follow-up, to detect tumor progression, to monitor response to therapy, and therefore to guide treatment plans.

## Materials and Methods

### Patient cohort and sample collection

All samples used in this study were previously analyzed as part of a prior study^7^, the details of which have been well documented. Written informed consent was obtained from 205 patients with primary cutaneous melanoma and 142 healthy control participants for this study^7^. Of the 205, 145 had American Joint Committee on Cancers (AJCC) clinical stages 0, I or II (i.e., early stage) and 60 patients had AJCC stages III or IV (i.e., late stage). The Human Research Ethics Committees of Edith Cowan University (No. 2932) and Sir Charles Gairdner Hospital (No. 2007-123) approved the study. Patients were recruited from the Medical Oncology Department of Sir Charles Gairdner Hospital and the Perth Melanoma Clinic at Hollywood Hospital (Perth, Western Australia), while the aged-matched, healthy group were recruited from the general population. Detailed patient cohort information is listed in **Table 1**. As reported previously ^7^, 2.5 ml of whole blood was collected into a PAXgene Blood RNA Tube (PreAnalytiX, Hombrechtikon, Switzerland) containing RNA stabilizers. Total RNA was isolated using the PAXgene Blood RNA Kit (Qiagen, Hilden, Germany) and then reverse transcription was carried by Omniscript Reverse Transcriptase (Qiagen). Clinical data and post-blood collection follow-up was collected for all patients over 3 y and 9 mon.

### Melanoma cell lines, RNA isolation, standard curve and quantitative PCR (qPCR)

Human melanoma cell line A375 (obtained from ATCC) were grown in Dubecco’s modified Eagle’s medium (DMEM; Gibco) media containing 10% fetal bovine serum (Hyclone) in T75 cm2 flask (Corning) until cells were ~70% confluent. Cells were then collected and total RNA was isolated using an RNeasy kit (QIAGEN) as per the manufacturers’ protocol. First-strand cDNA was synthesized using Superscript II reverse transcriptase (Invitrogen), followed by ribonuclease H treatment (Invitrogen) prior to performing quantitative real-time PCR. A standard curve, to quantify mRNA copy number, was constructed using larger PCR products (~500 bp) that included the target sequence used in qPCR. Reactions omitting reverse transcriptase were performed in each experiment as negative controls. Reactions were then run on a real-time PCR system (ABI StepOnePlus; Applied Biosystems). Gene expression of *Daple-V1, Daple-V2* and *S100A4* was detected with Taqman assay (Invitrogen), and relative gene expression was determined by normalizing to GAPDH using Relative standard curve method. Primer and probe sequences are as listed in **Supplementary Table 1** (**Table S1**).

### Analysis of melanoma-associated markers

These analyses were carried out as described previously ^7^. Briefly, total RNA (250 ng) was converted to cDNA (Omniscript Reverse Transcriptase, Qiagen), and included a no template control. qRT-PCR assays assessed the number of mRNA transcripts (level of expression) for three genes that have been previously shown to be associated with melanoma cells, and in some cases, indicative of stemness, i.e., *MLANA, ABCB5* and *MCAM. GAPDH* was assessed as an internal loading control because its levels are not upregulated in melanoma tissues or cultured cells relative to normal samples ^36^. These assays were carried out using SYBR GreenER qPCR SuperMix (Invitrogen) and 200 nmol L^−1^ of primer (**Table S1**) and an iCycler iQ5 Real-Time Thermocycler (Bio-Rad, Hercules, CA, U.S.A.) as per the manufacturer’s instructions. To prevent contamination, all PCR reactions contained uracil-N-glycosylase to prevent reamplification of carryover PCR products, and different steps were performed in separate ultraviolet-treatable areas. Melting point determination and gel electrophoresis confirmed the expected size and identity of PCR products. Every assay included a standard curve, negative controls (no template and reverse transcription control), positive control (A2058 cell line) and cDNA from a single healthy control.

### Statistical analyses

Statistical evaluation was performed using the GraphPad prism v5 software. Statistical significance of differences in marker expression among the patients with early *vs*. late stages of melanoma and normal healthy controls was analyzed using unpaired t-test. Optimal cut-off values for gene expression levels were derived by maximally selected log-rank statistics performed using R Software v2.15.0 ^37^. The relationship between high vs. low levels of gene expression and incidence of metastatic progression was investigated by Fisher’s exact test. To assess the relationship between gene expression and patient’s age, clinical stage of disease, Clark level, Breslow thickness and ulceration, we used parametric Pearson’s correlation and nonparametric Spearman’s correlation analyses. To estimate the correlation among the expression levels of various genes tested here, we used Correlation Matrix. To analyze the effect of high *vs*. low levels of gene expression and time-dependent survival probabilities we used Kaplan-Meier (KM) analyses. All statistical tests were performed two-sided, and *p*-values less than 0.05 were considered as statistically significant.

## DATA AVAILABILITY STATEMENT

All data generated or analyzed during this study are included in this published article (and its Supplementary Information files). The raw datasets generated during and/or analyzed during the current study are available from the corresponding author on reasonable request. The TCGA datasets analyzed during the current study are available in the cBioPortal repository, [http://www.cbioportal.org/].

## ACKNOWLEDGMENTS

This work was funded by NIH (R01CA160911 and R01CA100768) and a Translation and Clinical Research Award from the Moores Cancer Center to PG. J.E was supported by a NCI/NIH-funded Cancer Biology, Informatics & Omics (CBIO) Training Program (T32 CA067754) and a Postdoctoral Fellowship from the American Cancer Society (PF-18-101-01-CSM). Other funding support to MZ includes: NHMRC application number 1013349; Cancer and Palliative Care Research and Evaluation Unit WAPCN Small Grants 2010/11; and Cancer Council of WA Research Grants (to M.Z).

## AUTHOR CONTRIBUTIONS

Y.D. and P.G participated in research design; Y.D conducted experiments and performed statistical analysis. J.E. carried out bioinformatic analyses. A.L and M.Z provided all patient samples, and access to previously published data on melanoma-associated markers and provided critical guidance in the formulation of study design and manuscript reviews. Y.D, J.E., N.A and P.G. wrote the manuscript. P.G. conceived and supervised the project.

## REFERENCES

1 Faries, M. B., Steen, S., Ye, X., Sim, M. & Morton, D. L. Late recurrence in melanoma: clinical implications of lost dormancy. J Am Coll Surg 217, 27–34; discussion 34-26, doi:10.1016/j.jamcollsurg.2013.03.007 (2013).

2 Aguirre-Ghiso, J. A. Models, mechanisms and clinical evidence for cancer dormancy. Nature reviews. Cancer 7, 834–846, doi:10.1038/nrc2256 (2007).

3 Mocellin, S., Hoon, D., Ambrosi, A., Nitti, D. & Rossi, C. R. The prognostic value of circulating tumor cells in patients with melanoma: a systematic review and metaanalysis. Clinical cancer research: an official journal of the American Association for Cancer Research 12, 4605–4613, doi:10.1158/1078-0432.CCR-06-0823 (2006).

4 Ashworth, T. R. A case of cancer in which cells similar to those in the tumours were seen in the blood after death. Australian Medical Journal 14, 146–147 (1869).

5 Roland, C. L. et al. Detection of circulating melanoma cells in the blood of melanoma patients: a preliminary study. Melanoma research 25, 335–341, doi:10.1097/CMR.0000000000000168 (2015).

6 Freeman, J. B., Gray, E. S., Millward, M., Pearce, R. & Ziman, M. Evaluation of a multimarker immunomagnetic enrichment assay for the quantification of circulating melanoma cells. Journal of translational medicine 10, 192, doi:10.1186/1479-5876-10-192 (2012).

7 Reid, A. L. et al. Markers of circulating tumour cells in the peripheral blood of patients with melanoma correlate with disease recurrence and progression. The British journal of dermatology 168, 85–92, doi:10.1111/bjd.12057 (2013).

8 Koyanagi, K. et al. Multimarker quantitative real-time PCR detection of circulating melanoma cells in peripheral blood: relation to disease stage in melanoma patients. Clinical chemistry 51, 981–988, doi:10.1373/clinchem.2004.045096 (2005).

9 Khoja, L., Lorigan, P., Dive, C., Keilholz, U. & Fusi, A. Circulating tumour cells as tumour biomarkers in melanoma: detection methods and clinical relevance. Annals of oncology: official journal of the European Society for Medical Oncology / ESMO 26, 33–39, doi :10.1093/annonc/mdu207 (2015).

10 Bouwhuis, M. G. et al. Prognostic value of serial blood S100B determinations in stage IIB-III melanoma patients: a corollary study to EORTC trial 18952. European journal of cancer 47, 361–368, doi:10.1016/j.ejca.2010.10.005 (2011).

11 Carrillo, E. et al. Prognostic value of RT-PCR tyrosinase detection in peripheral blood of melanoma patients. Disease markers 22, 175–181 (2006).

12 Steen, S., Nemunaitis, J., Fisher, T. & Kuhn, J. Circulating tumor cells in melanoma: a review of the literature and description of a novel technique. Proc (Bayl Univ Med Cent) 21, 127–132 (2008).

13 Hoon, D. S. et al. Molecular markers in blood as surrogate prognostic indicators of melanoma recurrence. Cancer research 60, 2253–2257 (2000).

14 Gogas, H. et al. Prognostic significance of the sequential detection of circulating melanoma cells by RT-PCR in high-risk melanoma patients receiving adjuvant interferon. British journal of cancer 87, 181–186, doi:10.1038/sj.bjc.6600419 (2002).

15 Shah, K. V., Chien, A. J., Yee, C. & Moon, R. T. CTLA-4 is a direct target of Wnt/beta-catenin signaling and is expressed in human melanoma tumors. The Journal of investigative dermatology 128, 2870–2879, doi:10.1038/jid.2008.170 (2008).

16 Anastas, J. N. et al. WNT5A enhances resistance of melanoma cells to targeted BRAF inhibitors. The Journal of clinical investigation 124, 2877–2890, doi:10.1172/JCI70156 (2014).

17 Atkinson, J. M. et al. Activating the Wnt/beta-Catenin Pathway for the Treatment of Melanoma--Application of LY2090314, a Novel Selective Inhibitor of Glycogen Synthase Kinase-3. PloS one 10, e0125028, doi:10.1371/journal.pone.0125028 (2015).

18 Aznar, N. et al. Daple is a novel non-receptor GEF required for trimeric G protein activation in Wnt signaling. Elife 4, e07091, doi:10.7554/eLife.07091 (2015).

19 Aznar,N. et al. A Daple-Akt feed forward loop enhances non-canonical Wnt signals by compartmentalizing beta-Catenin. Molecular biology of the cell, doi:10.1091/mbc.E17-06-0405 (2017).

20 Aznar, N. et al. Convergence of Wnt, growth factor, and heterotrimeric G protein signals on the guanine nucleotide exchange factor Daple. Sci Signal 11, doi:10.1126/scisignal.aao4220 (2018).

21 Ara, H. et al. Role for Daple in non-canonical Wnt signaling during gastric cancer invasion and metastasis. Cancer science 107, 133–139, doi:10.1111/cas.12848 (2016).

22 Dunkel, Y. et al. Two Isoforms of the Guanine Nucleotide Exchange Factor, Daple Cooperate as Tumor Suppressors. FASEB journal: official publication of the Federation of American Societies for Experimental Biology (2018).

23 Barbazan, J. et al. Prognostic Impact of Modulators of G proteins in Circulating Tumor Cells from Patients with Metastatic Colorectal Cancer. Sci Rep 6, 22112, doi:10.1038/srep22112 (2016).

24 Cardenas-Navia, L. I. et al. Novel somatic mutations in heterotrimeric G proteins in melanoma. Cancer biology & therapy 10, 33–37 (2010).

25 O’Hayre, M. et al. The emerging mutational landscape of G proteins and G-protein-coupled receptors in cancer. Nat Rev Cancer 13, 412–424, doi:10.1038/nrc3521 (2013).

26 Van Raamsdonk, C. D. et al. Frequent somatic mutations of GNAQ in uveal melanoma and blue naevi. Nature 457, 599–602, doi:10.1038/nature07586 (2009).

27 Smith, B. et al. Detection of melanoma cells in peripheral blood by means of reverse transcriptase and polymerase chain reaction. Lancet 338, 1227–1229 (1991).

28 Bettum, I. J. et al. Metastasis-associated protein S100A4 induces a network of inflammatory cytokines that activate stromal cells to acquire pro-tumorigenic properties. Cancer letters 344, 28–39, doi:10.1016/j.canlet.2013.10.036 (2014).

29 Dahlmann, M., Kobelt, D., Walther, W., Mudduluru, G. & Stein, U. S100A4 in Cancer Metastasis: Wnt Signaling-Driven Interventions for Metastasis Restriction. Cancers 8, doi:10.3390/cancers8060059 (2016).

30 Maelandsmo, G. M. et al. Differential expression patterns of S100A2, S100A4 and S100A6 during progression of human malignant melanoma. International journal of cancer 74, 464–469 (1997).

31 Luo, X. et al. Methylation-mediated loss of SFRP2 enhances melanoma cell invasion via Wnt signaling. Am J Transl Res 8, 1502–1509 (2016).

32 De Giorgi, V. et al. Circulating benign nevus cells detected by ISET technique: warning for melanoma molecular diagnosis. Archives of dermatology 146, 1120–1124, doi:10.1001/archdermatol.2010.264 (2010).

33 Clawson, G. A. et al. Circulating tumor cells in melanoma patients. PLoS One 7, e41052, doi:10.1371/journal.pone.0041052 (2012).

34 Lee, J. et al. Conditions for the differentiation of melanocyte-precursor cells from human cord blood-derived mesenchymal stem cells. African J Biotechnology 9: 5975–5977 (2010).

35 Ohta, S. et al. Generation of human melanocytes from induced pluripotent stem cells. PloS one 6, e16182, doi:10.1371/journal.pone.0016182 (2011).

36 Giricz, O., Lauer-Fields, J. L. & Fields, G. B. The normalization of gene expression data in melanoma: investigating the use of glyceraldehyde 3-phosphate dehydrogenase and 18S ribosomal RNA as internal reference genes for quantitative real-time PCR. Anal Biochem 380, 137–139, doi:10.1016/j.ab.2008.05.024 (2008).

37 Budczies, J. et al. Cutoff Finder: a comprehensive and straightforward Web application enabling rapid biomarker cutoff optimization. PLoS One 7, e51862, doi:10.1371/journal.pone.0051862 (2012).

